# Natural genetic variation in the pheromone production of *C. elegans*

**DOI:** 10.1101/2022.11.23.517769

**Authors:** Daehan Lee, Bennett W. Fox, Diana C. F. Palomino, Oishika Panda, Francisco J. Tenjo, Emily J. Koury, Kathryn S. Evans, Lewis Stevens, Pedro R. Rodrigues, Aiden R. Kolodziej, Frank C. Schroeder, Erik C. Andersen

## Abstract

From bacterial quorum sensing to human language, communication is essential for social interactions. Nematodes produce and sense pheromones to communicate among individuals and respond to environmental changes. These signals are encoded by different types and mixtures of ascarosides, whose modular structures further enhance the diversity of this nematode pheromone language. Interspecific and intraspecific differences in this ascaroside pheromone language have been described previously, but the genetic basis and molecular mechanisms underlying the variation remain largely unknown. Here, we analyzed natural variation in the production of 44 ascarosides across 95 wild *Caenorhabditis elegans* strains using high-performance liquid chromatography coupled to high-resolution mass spectrometry (HPLC-HRMS). By cross-analyzing genomes and *exo*-metabolomes of wild strains, we discovered quantitative trait loci (QTL) that underlie the natural differences in pheromone bouquet composition. Fine mapping of the QTL further uncovered associations between mitochondrial metabolism and pheromone production. Our findings demonstrate how natural genetic variation in core metabolic pathways can affect the production of social signals.

## Introduction

“Pheromone” is an informative chemical or mixture of chemicals that an organism produces and secretes into the environment, affecting the behavior, physiology, and development of other individuals. Nematodes use pheromones called ascarosides^1,2^, which consist of the dideoxy sugar, ascarylose, linked to diverse fatty acid (FA) side chains as well as to derivatives of amino acids, folate, and other primary metabolites^3^ (Fig. 1a). The nematode sensory system perceives distinct combinations and concentrations of ascarosides^4^, which in turn modulate a variety of biological processes, including developmental plasticity, social and sexual behaviors, olfactory learning, stress response, reproduction, and longevity^1,5–11^.

**Fig. 1.**
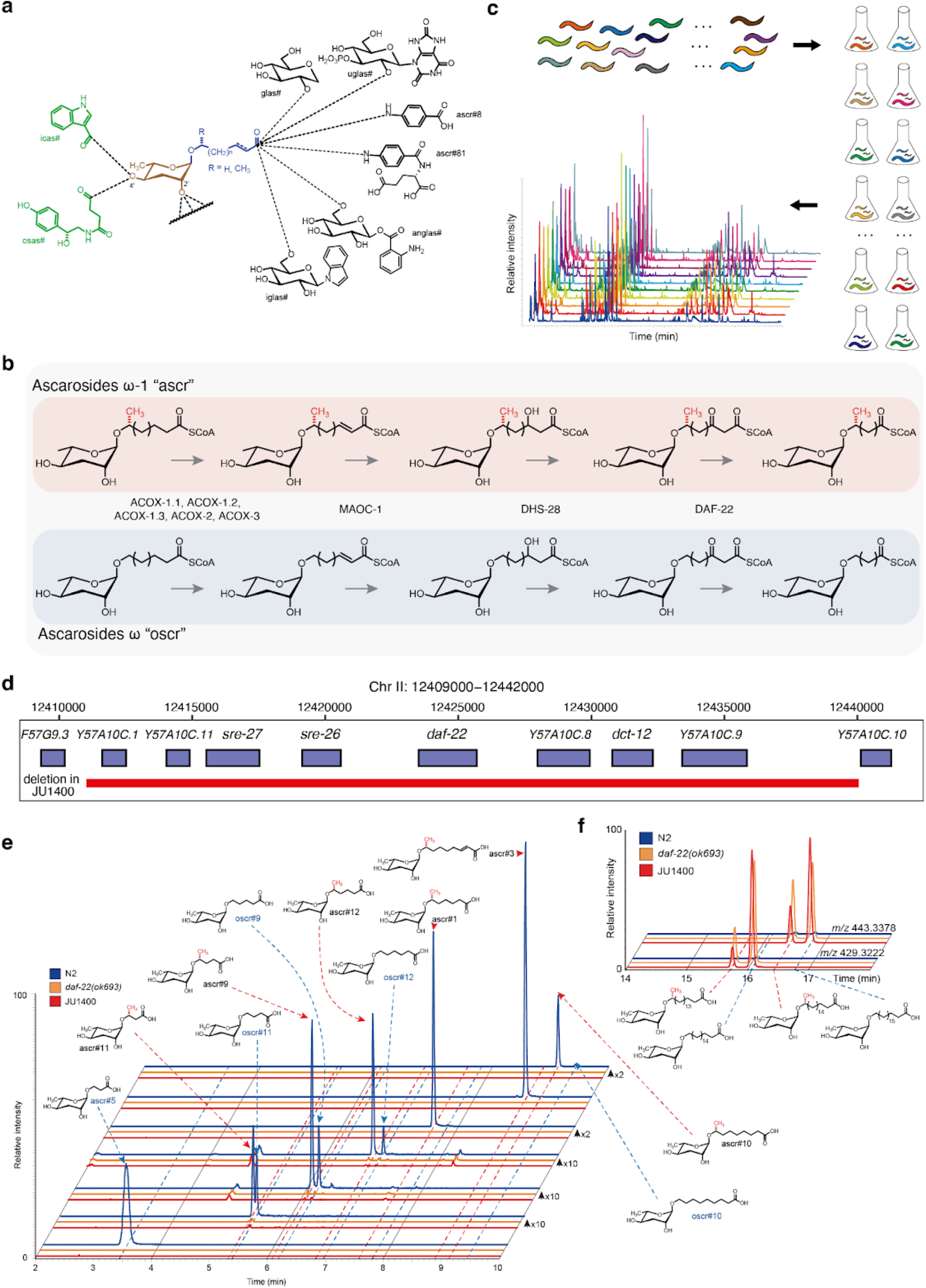
A natural deletion in *daf-22* gene was found in pheromone deficient wild *C. elegans* strain. (a) Chemical structures of the ascarosides. Simple ascarosides consist of the dideoxy sugar ascarylose (brown) and a FA-like side chain of varying lengths (blue). Modular ascarosides incorporate additional building blocks from other primary metabolic pathways, e.g., indole 3-carboxylic acid (icas#, green) derived from tryptophan, the neurotransmitter octopamine (osas#, green), and a variety of C-terminal modified ascarosides, including folate derivatives (ascr#8, ascr#81) and several glucosides (black)^30^. (b) Chain-shortening of ascarosides by enzymes in the peroxisomal β-oxidation pathway. Ascarosides of the two classes (“ascr” and “oscr”) are derived from very long-chain precursors. (c) Schematic of the ascaroside profiling experiments, in which 95 *C. elegans* strains were grown in liquid cultures, and the conditioned media extracted and analyzed by HPLC-HRMS, then correlated with genomic data to identify QTL underlying natural differences in pheromone production. (d) A 29,011 bp natural deletion in the JU1400 strain encompassing *daf-22* and seven additional genes. (e) Extracted ion chromatograms (EICs) corresponding to several short- and medium-chain ascarosides, as indicated, in the N2 strain (WT), *daf-22(ok693)*, and the JU1400 strain *exo*-metabolome extracts from synchronized adults. Y-axes are scaled as indicated to clearly show lower intensity metabolites. (f) EICs for *m/z* 429.3222 and 443.3378, corresponding to precursor ascarosides with C_18_ and C_19_ sidechains, respectively, in the N2 strain (WT), *daf-22(ok693)*, and the JU1400 strain *exo*-metabolome extracts from synchronized adults.

Studies of the model nematode species *Caenorhabditis elegans* have uncovered several ascaroside biosynthetic pathways. One of the key biochemical reactions in ascaroside production is the iterative shortening of the fatty acid (FA) side chain by peroxisomal β-oxidation (Fig. 1b). Mutations in peroxisomal β-oxidation genes impair the production of functional short- and medium-chained ascarosides that control development and behavior^12–16^. Analysis of peroxisomal β-oxidation mutants, in particular *daf-22(ok693)*, revealed an accumulation of long-chained precursors with both odd and even numbers of carbons in the side chain^15^. Although the roles of many genes involved in the production of ascarosides have been characterized (*e*.*g*., *daf-22, dhs-28, maoc-1*, and acyl-CoA oxidase (ACOX) orthologs)^15,17–20^, the upstream pathway that produces long-chained precursor ascarosides is largely unknown.

Ascaroside pheromones are a universal nematode chemical language found across diverse parasitic and free-living species2, but the repertoire of ascaroside pheromones varies from species to species. For example, a dimeric ascaroside discovered in *Pristionchus pacificus*, dasc#1, regulates the mouth-form dimorphism underlying its facultative predatory lifestyle^21^. In addition, intraspecific quantitative variation in pheromone production has been observed in both *C. elegans* and *P. pacificus* species^2,22,23^. These discoveries suggest that ascaroside biosynthetic pathways vary within and across species. In line with the natural variation in pheromone production, natural differences in pheromone responses have been demonstrated as well^22,24–27^. Here, we characterized the genetic basis of natural variation in *C. elegans* pheromone production. We analyzed the secreted metabolites from 95 wild *C. elegans* strains and profiled their pheromone bouquets by measuring relative abundances of 44 different ascarosides. Our quantitative genetic analysis of heritable variation in ascaroside production revealed a novel link between natural differences in metabolism and chemical communication of the species.

## Results

### A peroxisomal β-oxidation gene is deleted in a pheromone-less wild *C. elegans* strain

To investigate the intraspecific variation in *C. elegans* pheromone production, we analyzed the *exo*-metabolomes of 95 wild strains using high-performance liquid chromatography coupled to high-resolution mass spectrometry (HPLC-HRMS) (Fig. 1c, Methods). Because ascaroside biosynthesis is affected by diverse factors including sex, developmental timing, and nutrition^28^, we chose to analyze the *exo*-metabolomes of synchronized hermaphrodites at the young adult stage. Among thousands of metabolites, we identified and quantified 44 ascarosides (Methods, Supplementary Fig. 1). Notably, short- and medium-chain ascarosides were severely depleted in a single wild strain (JU1400), suggesting that this strain might have an impaired ascaroside biosynthetic pathway. Previously, mutations in peroxisomal β-oxidation genes (*e*.*g*., *daf-22, dhs-28, maoc-1*) (Fig. 1b) were shown to abolish the production of short- and medium-chain ascarosides^15^. To investigate whether the JU1400 strain has an impaired peroxisomal β-oxidation pathway, we examined sequences of these genes and identified a large deletion (29 kb) in the *daf-22* locus, which completely removes *daf-22* and seven neighboring genes (Fig. 1d). The JU1400 strain displayed a similar phenotype as a *daf-22* loss-of-function mutant in which short- and medium-chain ascarosides were absent but long-chain precursors were present (Fig. 1e, f). Intriguingly, the ratio of very long-chain (ω-1)- to ω-ascarosides in *daf-22(ok693)* and JU1400 was similar (Fig. 1f), suggesting that the ratio of these precursors is controlled independently of peroxisomal β-oxidation. We extended our analysis of the *daf-22* locus to 538 wild genomes^29^ but did not find the same deletion nor any nonsense mutation in other wild strains, suggesting that the loss of a β-oxidation gene that leads to the severe impairment of ascaroside production is rare in the natural *C. elegans* population.

### The pheromone bouquet varies among wild *C. elegans* strains

*C. elegans* produces and releases a diverse collection of ascarosides with different lengths of FA side chains as well as other types of modifications (Fig. 1a). We compared the composition and abundances of pheromones among 94 wild *C. elegans* strains (Fig. 2a,b, Supplementary Data 1), excluding the JU1400 strain that lacks the majority of short- and medium-chain ascarosides. For each strain, we calculated the intensity of each ascaroside relative to the sum of the 44 measured ascarosides (henceforth, referred to as relative abundance). On average, we found that two major ascarosides, ascr#5 and ascr#3, which are derived from the ω and ω-1 pathways, comprised 51.2% and 21.4% of measured ascarosides, respectively (Fig. 2c). The rest of the identified ascarosides comprised 0.004% to 7.4% and the relative abundances of each ascaroside varied from strain to strain, though to a different extent (Fig. 2b,c, Supplementary Fig. 2). For example, a pheromone signal that promotes dauer formation at high concentration and aggregation at picomolar concentrations, referred to as icas#9^8,31^, was not detected in the ECA36 strain but it comprised 3.3% of the total ascarosides in the CB4856 strain (Fig. 2d). Notably, ECA36 possesses a nonsense mutation in *cest-3*, which encodes the enzyme required for 4’ attachment of indole 3-carboxylic acid to the ascarylose core^32^. By contrast, ascr#11 was much less variable than icas#9 across the 94 wild strains, as its relative abundance ranged from 2.1% to 5% (Fig. 2e).

**Fig. 2.**
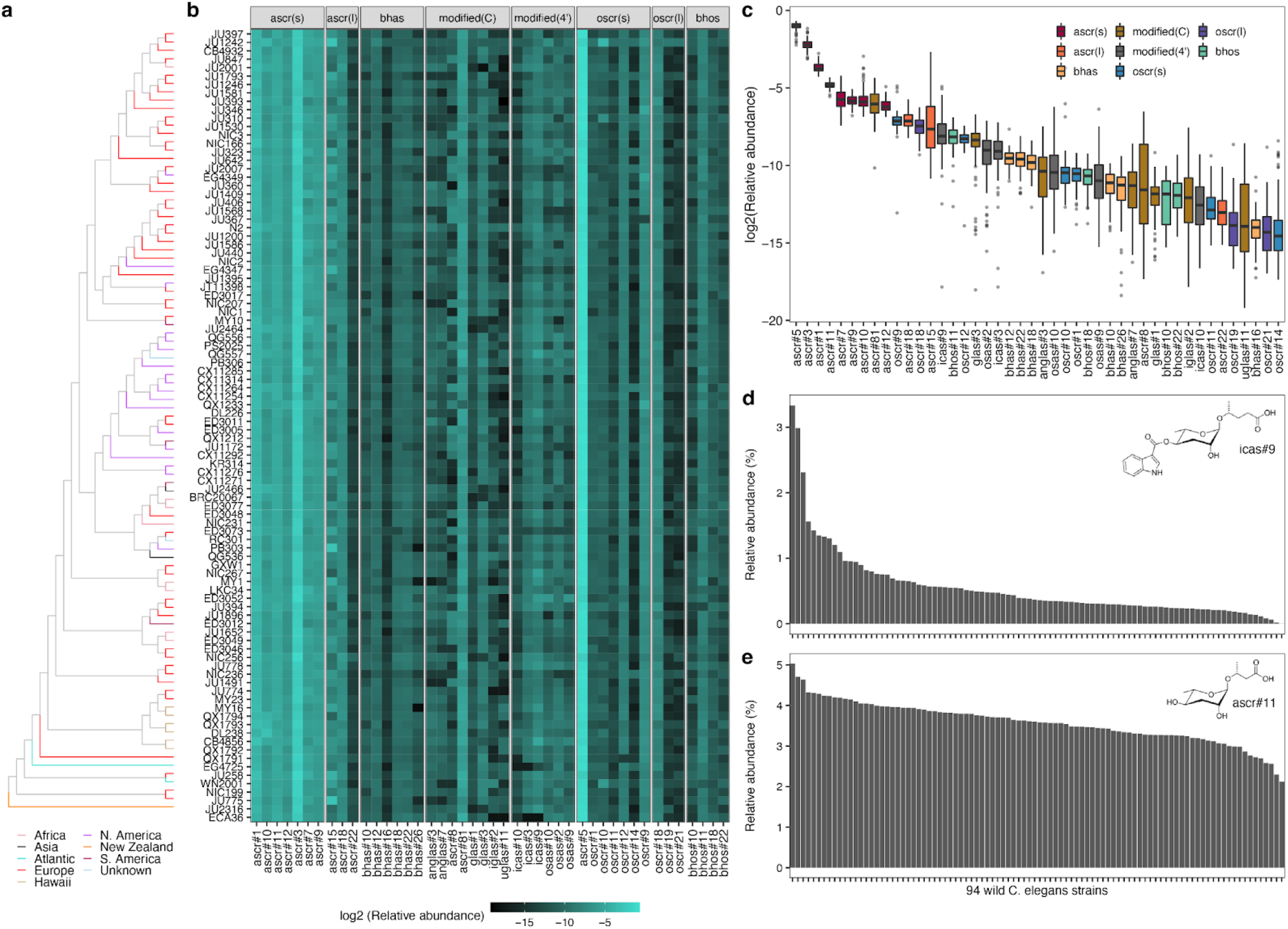
Pheromone bouquet varies across wild *C. elegans* strains. (a) Neighbor-joining tree of 94 wild *C. elegans* strains analyzed in this study (except the JU1400 strain) generated from 963,027 biallelic segregating sites. The terminal branch is colored by the strain’s geographic origin. (b) A heatmap showing relative abundances of 44 ascarosides. Each row represents one of the 94 wild strains, ordered by genome-wide relatedness. Ascarosides are grouped by structural similarity. (c) Tukey box plots of the relative abundances of 44 ascarosides across 94 wild *C. elegans* strains are shown with outlier strain data points plotted. The horizontal line in the middle of the box is the median, and the box denotes the 25th to 75th quantiles of the data. The vertical line represents the 1.5x interquartile range. Box plots are colored by structural properties that were used for ascaroside grouping in (b). (d,e) Bar plots showing relative abundances of icas#9 (d) and ascr#11 (e) across 94 wild *C. elegans* strains, ordered by relative abundance.

To investigate the genetic contributions to the natural variation in ascaroside abundances, we analyzed the heritabilities for relative abundance traits of 44 ascarosides. We found that the narrow-sense heritabilities (*h*^*2*^) of 44 ascaroside abundance traits ranged from 0% to 80% (Fig. 3a). Variation in the icas#9 abundance trait exhibited the highest heritability (80%), followed by ascr#10 (67%), icas#10 (59%), and ascr#5 (57%). All indole-3-carboxylic acid ascarosides (icas#) showed high heritabilities (>50%), whereas differences in β-hydroxylated ω-ascarosides (bhos#10, bhos#11, bhos#18, bhos#22) were not explained by additive genetic factors (*h*^*2*^=0%). To focus on genetic differences in pheromone production, we chose 23 ascarosides that showed at least 10% of total additive genetic variance for the following analyses.

**Fig. 3.**
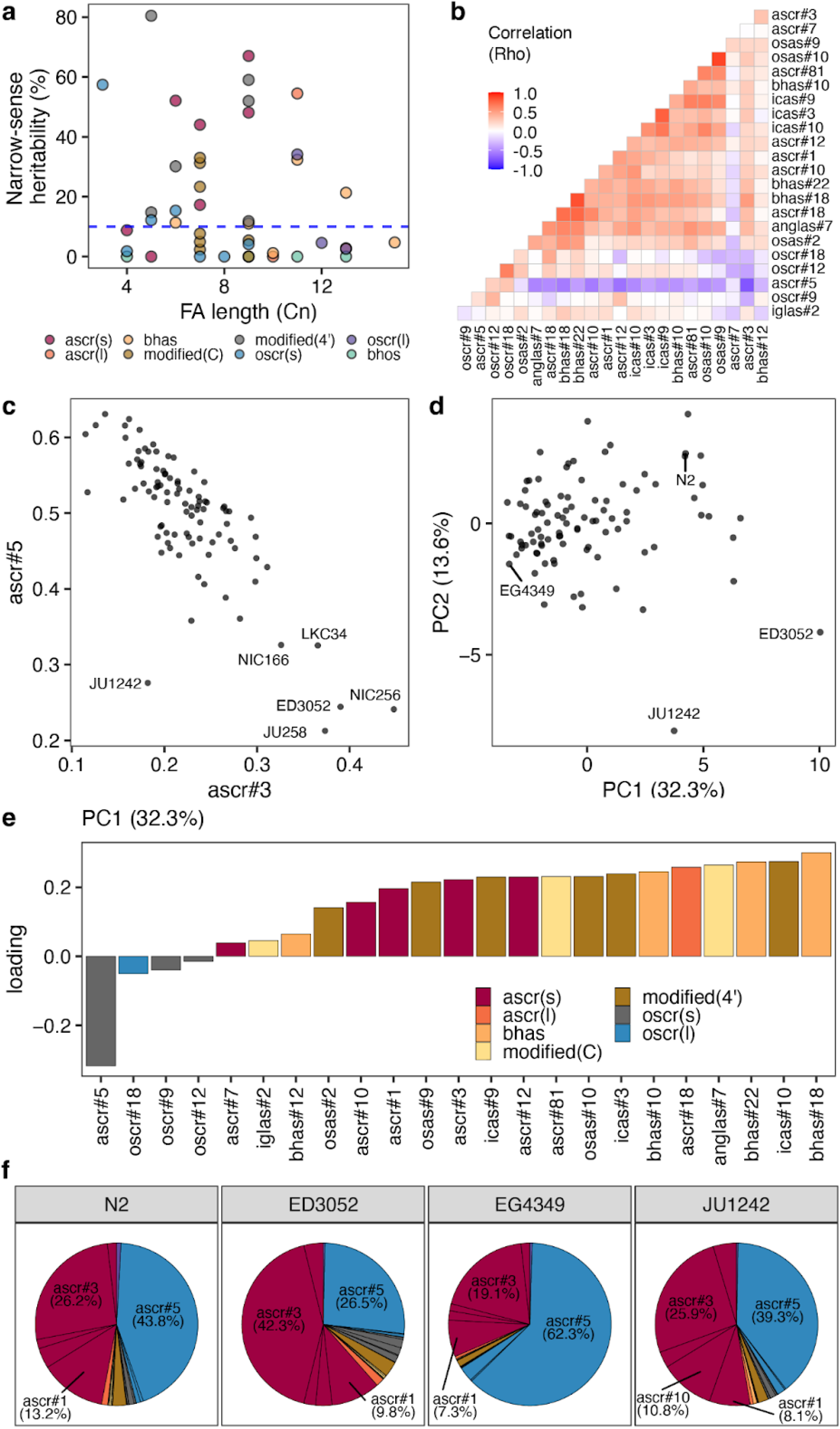
Relative abundances of ω- and (ω-1)-ascarosides are inversely correlated. (a) Narrow-sense heritabilities of 44 different ascarosides are shown. Each point corresponds to one of the 44 ascarosides and is colored by the structural property. The blue dashed horizontal line shows the heritability threshold (10%) for downstream analyses. The length of the FA side chain is shown on the x-axis. (b) A heatmap showing correlation (Spearman’s rho) of relative abundance traits between two different ascarosides. (c) A scatter plot showing relative abundances of ascr#3 (x-axis) and ascr#5 (y-axis). Each point corresponds to one of the 94 wild *C. elegans* strains. (d) Plots of the 94 wild strains according to their values for each of the two significant axes of variation, as determined by PCA of the ascaroside pheromone bouquet. (e) Relative abundance trait loadings of the first PC that explains up to 32.3% of the total variance in the trait data are shown. (f) Pie charts showing the relative abundances of 23 ascarosides with high heritability (>10%) for four wild *C. elegans* strains labeled in (d) are shown. (e,f) Bar plots and pie charts are colored by the structural properties as in (a).

We analyzed the correlation patterns among these 23 heritable traits and found that negative correlations are prevalent between ω-ascarosides and (ω-1)-ascarosides. For example, the most abundant ω-ascaroside (ascr#5) showed negative correlations with all (ω-1)-ascarosides but positive correlations with all ω-ascarosides (oscr#9, oscr#12, oscr#18) (Fig. 3b). The strongest correlations (Spearman’s *rho* = -0.81) were observed between ascr#5 and the most abundant (ω-1)-ascaroside (ascr#3) (Fig. 3b,c). Furthermore, the ratio between ascr#3 and ascr#5 was remarkably heritable (*h*^*2*^ = 82.1%), which was even higher than the *h*^*2*^ values of the relative abundance traits for ascr#3 (48.1%) or ascr#5 (57.4%). We observed a similar trend using principal component analysis (PCA) of the 23 heritable traits (Fig. 3d-f, Methods). The first principal component (PC) that explained 32.3% of the variance in the dataset had negative loadings for all ω-ascarosides (ascr#5, oscr#9, oscr#12, oscr#18) and positive loadings for all (ω-1)-ascarosides (Fig. 3e). Because ω- and (ω-1)-ascarosides are derived from parallel β-oxidation of very-long chain precursors^28^, these results suggest that the relative amounts of ω- and (ω-1)-starting materials vary across the species.

### A common genomic locus underlies variation in ascaroside biosynthesis

To characterize genomic loci underlying observed natural differences in the composition of the pheromone bouquet, we performed genome-wide association (GWA) mappings (Supplementary Table 1). We identified four quantitative trait loci (QTL) from the mapping of ascr#3:ascr#5 ratio trait (Fig. 4a,b), including the most significant genomic region on the right arm of chromosome II (peak marker at II:13,692,928), which explained 71.8% of the phenotypic variance. Specifically, five wild strains (ED3052, JU258, LKC34, NIC166, and NIC256) had high ascr#3:ascr#5 ratios (≥ 1) and the non-reference (ALT) allele at the peak position (Fig. 4b). We also performed GWA mapping for heritable (>10%) relative abundance traits of 23 ascarosides and identified quantitative trait loci (QTL) from 20 traits (Fig. 4c). The ascr#3:ascr#5 trait QTL overlapped with QTL that were mapped for relative abundance traits; ChrIIR-QTL overlapped with 11 ascarosides (anglas#7, ascr#1, ascr#3, ascr#5, ascr#7, ascr#12, ascr#81, bhas#18, bhas#22, osas#2, and oscr#12) and three other QTL (ChrIIL-QTL, ChrIV-QTL, and ChrX-QTL) also overlapped with QTL from other relative abundance traits (Fig. 4c). These results together suggest that shared genomic loci (“QTL hotspots”) harbor variants with a broad impact on ascaroside biosynthesis.

**Fig. 4.**
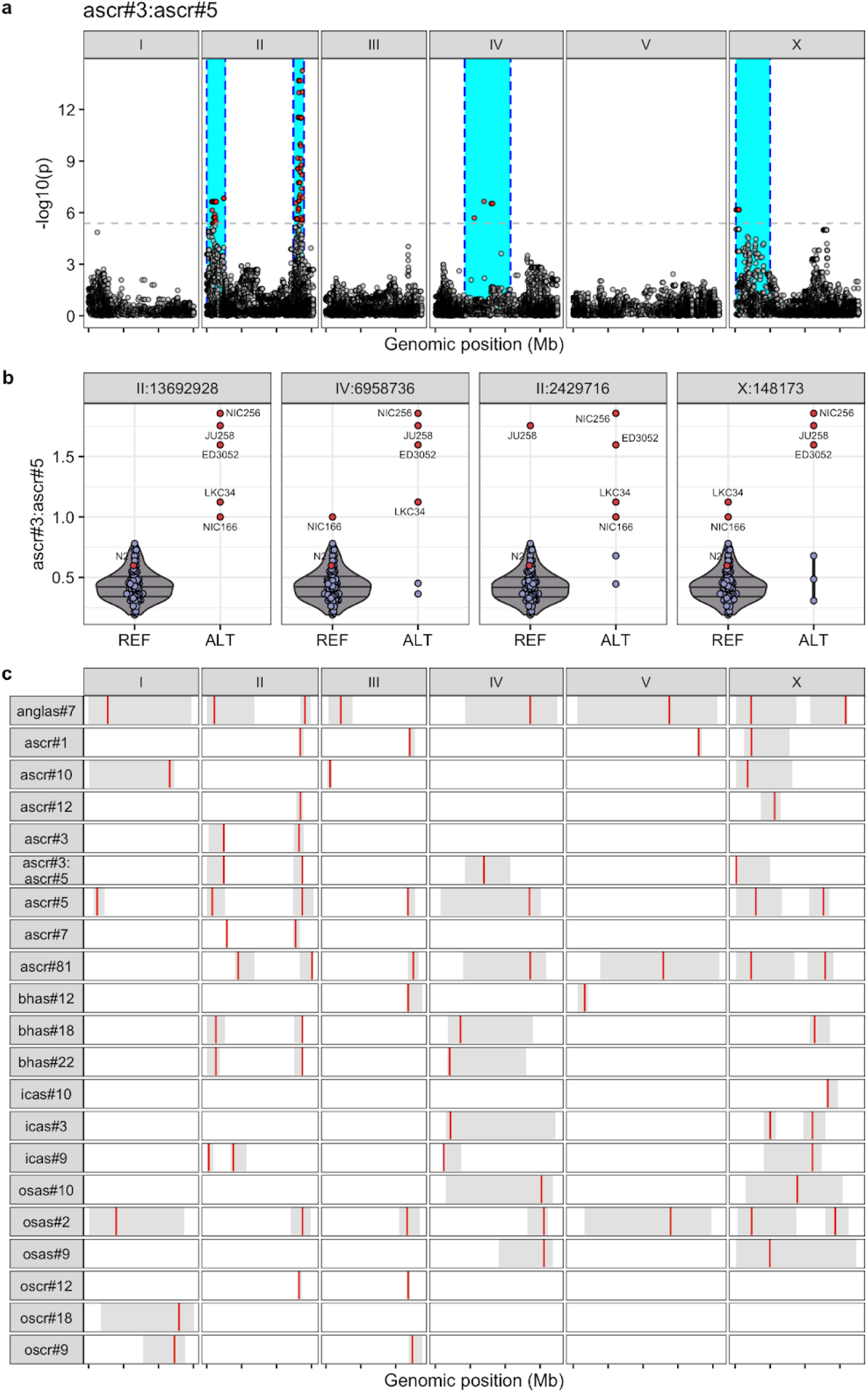
QTL hotspots underlie natural variation in various ascarosides. (a) A Manhattan plot for GWA mapping of the ascr#3:ascr#5 ratio trait is shown. Each dot represents a single-nucleotide variant (SNV) that is present in at least 5% of the 94 wild strains. The genomic position in Mb, separated by chromosome, is plotted on the x-axis, and the statistical significance of the correlation between genotype and phenotype is plotted on the y-axis. The dashed horizontal line denotes the Bonferroni-corrected p-value threshold using independent markers correcting for LD (genome-wide eigen-decomposition significance threshold). SNVs are colored red if they pass this threshold. The region of interest for each QTL is represented by a cyan rectangle. (b) Beeswarm plots of phenotypes split by peak marker position of the four QTL from (a). Each dot corresponds to the phenotype of an individual strain, which is plotted on the y-axis. Strains are grouped by their genotype at each peak QTL position. Dots for reference N2 strain and five high trait value strains are colored red. (c) A summary plot showing the GWA mapping results including location and range of QTL for relative abundance traits for 20 ascarosides with high trait heritability (>10%) and the ascr#3:ascr#5 ratio trait. Each red bar corresponds to the peak position of the QTL and each grey box represents the region of interest for each QTL.

We focused on the QTL hotspot on the right arm of chromosome II (ChrIIR-QTL), which explained the largest fraction of variance for the ascr#3:ascr#5 ratio trait and also was mapped for the largest number of relative abundance traits. To identify a quantitative trait gene (QTG) underlying this hotspot, we performed a fine-mapping for the ascr#3:ascr#5 ratio trait (Fig. 5a). Among genetic variants that were predicted to impact gene function (*e*.*g*., missense, frameshift), one of the most significantly associated variants (-log_10_p=5.71) was in the *mecr-1* gene, which encodes a mitochondrial trans-2-enoyl-CoA reductase. We found that of the five wild strains with high ascr#3:ascr#5 ratios, four strains (ED3052, JU258, LKC34, and NIC256) harbored the G159V missense variant in *mecr-1* (Fig. 5b), suggesting that this allele could cause increased production of ascr#3, a reduction of ascr#5, or both. This gene encodes a key enzyme in the mitochondrial FA synthesis (mtFAS) pathway^33^, whose potential interactions with ascaroside biosynthesis have not been described previously.

**Fig. 5.**
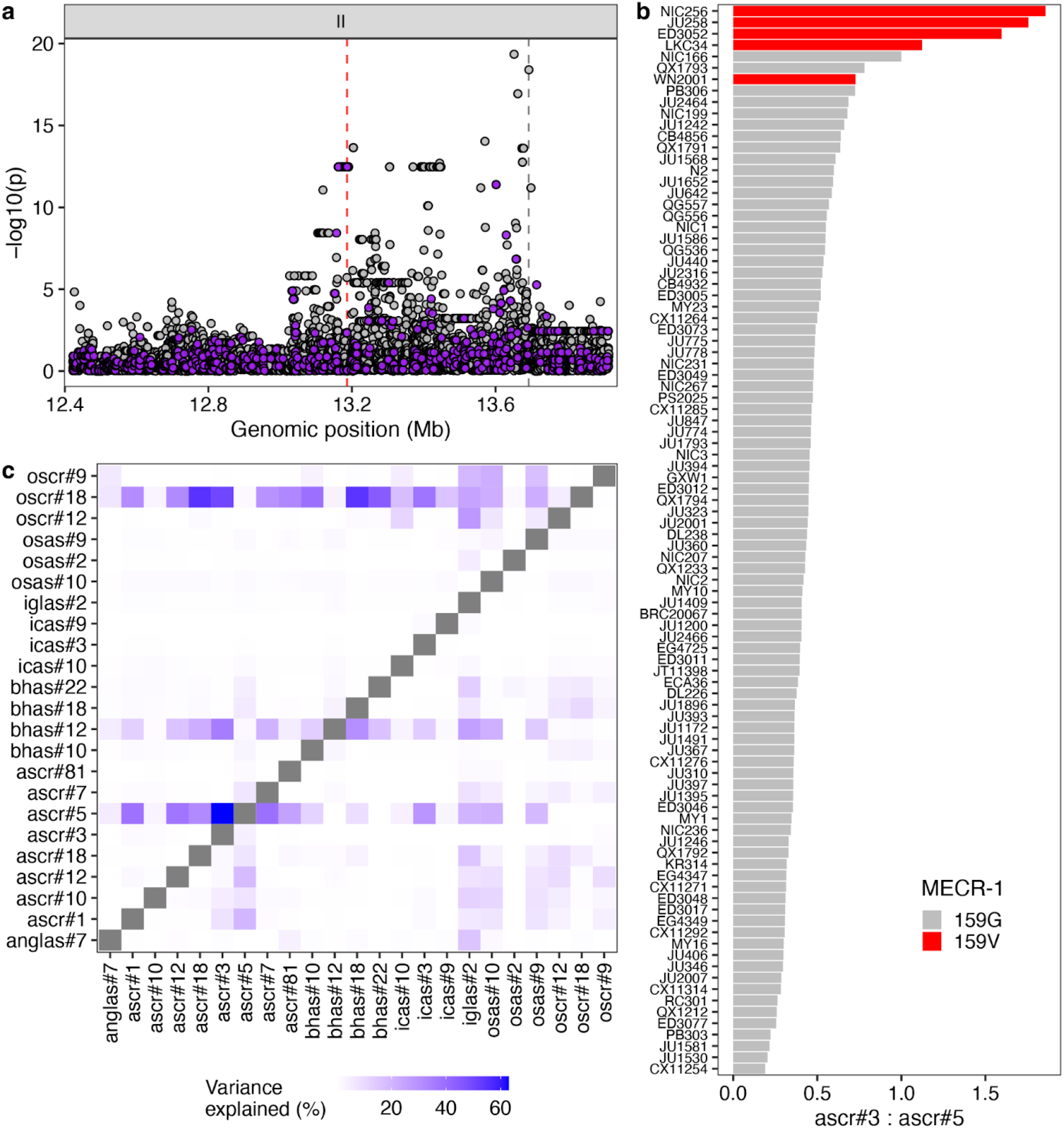
*mecr-1* variant is associated with natural differences in ascaroside production. (a) Fine mapping of the ChrIIR-QTL (II:12,422,412-13,692,928) for the ascr#3:ascr#5 ratio trait is shown. Each dot represents an SNV that is present in at least 5% of the 94 wild strains. The association between the SNV and ascr#3:ascr#5 trait value is shown on the y-axis, and the genomic position of the SNV is shown on the x-axis. SNVs with high or moderate impact inferred from SnpEff are colored purple. (b) A bar plot for the ascr#3:ascr#5 trait value of 94 wild *C. elegans* strains. Wild strains with the MECR-1(G159V) variant are colored red. (c) A heatmap showing amounts of variance explained by MECR-1(G159V) variant for pairwise ratio traits of 23 ascarosides with high trait heritability.

We extended our analysis to the association between the *mecr-1* variant and pairwise ratio traits of all 23 analyzed ascarosides. We found that the MECR-1(G159V) variant explained much of the phenotypic variance for many traits (Fig. 5c). Specifically, the MECR-1(G159V) variant was frequently associated with differences between ω-ascarosides (ascr#5, oscr#9, oscr#12, oscr#18) and (ω-1)-ascarosides (*e*.*g*., ascr#1, ascr#3, ascr#7, bhas#18, icas#3, osas#2), suggesting that this variant could affect the balance between the two parallel ascaroside production pathways. In addition, the relative ratio traits between bhas#12, an (ω-1)-oxygenated β-hydroxy ascaroside, and many (ω-1)-ascarosides were associated with the MECR-1(G159V) variant. Relative abundances of iglas#2, osas#9, and osas#10 to many other ascarosides (both ω-ascarosides and (ω-1)-ascarosides) were also associated with this same variant. Taken together, these results suggested that the *mecr-1* variant might broadly affect ascaroside biosynthesis pathways.

### Natural genetic variants in the mtFAS pathway are associated with the variation in ascaroside biosynthesis

To test whether the MECR-1(G159V) variant underlies natural differences in ascaroside production, we performed allele-replacement experiments. Using CRISPR-Cas9 genome editing of *mecr-1*, we tried to substitute the glycine at position 159 with a valine in the N2 strain and the valine at position 159 with a glycine in the ED3052 strain (Methods). We could successfully substitute the glycine with valine in the N2 strain, but we failed to edit the valine with glycine in the ED3052 strain after extensive trials (n. of injected animals = 63). We also tried to introduce the same valine-to-glycine edit in the NIC256 strain (n. of injected animals = 43) but also failed. We concluded that glycine at this residue is not compatible in these genetic backgrounds, likely from uncharacterized genetic interactions with other alleles. Using allele-replacement strains in the N2 background, we examined the phenotypic effects of the MECR-1(G159V) variant. We found that two independent N2 MECR-1(G159V) lines did not recapitulate the association between the MECR-1(G159V) variant and the ascr#3:ascr#5 ratio trait (Supplementary Fig. 3).

Because the MECR-1(G159V) allele-replacement strains did not reproduce the QTL effect, we hypothesized that multiple linked alleles together might contribute to the QTL effect. Notably, ALT alleles at the peak markers of all four ascr#3:ascr#5 ratio QTL (II:2,429,716; II:13,692,928; IV:6,958,736, X:148,173) were associated with greater trait values (Fig. 4b), and these alleles displayed strong linkage disequilibrium (LD) (Supplementary Fig. 4). Specifically, two QTL that are on opposite arms of chromosome II (ChrIIL-QTL-ChrIIR-QTL) showed high LD (*r*^*2*^ = 0.509). Surprisingly, we detected the same level of LD between QTL on different chromosomes (ChrIIR-QTL-ChrV-QTL, Supplementary Fig. 4). These intrachromosomal and interchromosomal LD might reflect genetic interactions between MECR-1(G159V) and other uncharacterized alleles on pheromone production.

To test this hypothesis, we performed a fine-mapping for the ChrIIL-QTL, which has a much smaller confidence interval (2.6 Mb) than the ChrIV-QTL (6.5 Mb) that spans almost half of chromosome IV (Fig. 6a). Within the ChrIIL-QTL for the ascr#3:ascr#5 ratio trait, the most significantly associated variant predicted to impact gene function was in *pod-2*, an ortholog of human ACACA (acetyl-CoA carboxylase alpha), which acts upstream of *mecr-1* in the mtFAS pathway (Fig. 6b). The alternative POD-2(1516Y) allele is associated with a higher ascr#3:ascr#5 ratio than the reference POD-2(1516H) allele (Fig. 6c). Furthermore, the association patterns of pairwise ratio traits are similar to that of MECR-1(G159V) variant (Supplementary Fig. 5). Most importantly, three wild strains that exhibit extremely high ascr#3:ascr#5 ratios carry alternative alleles for both genes (Fig. 6c).

**Fig. 6.**
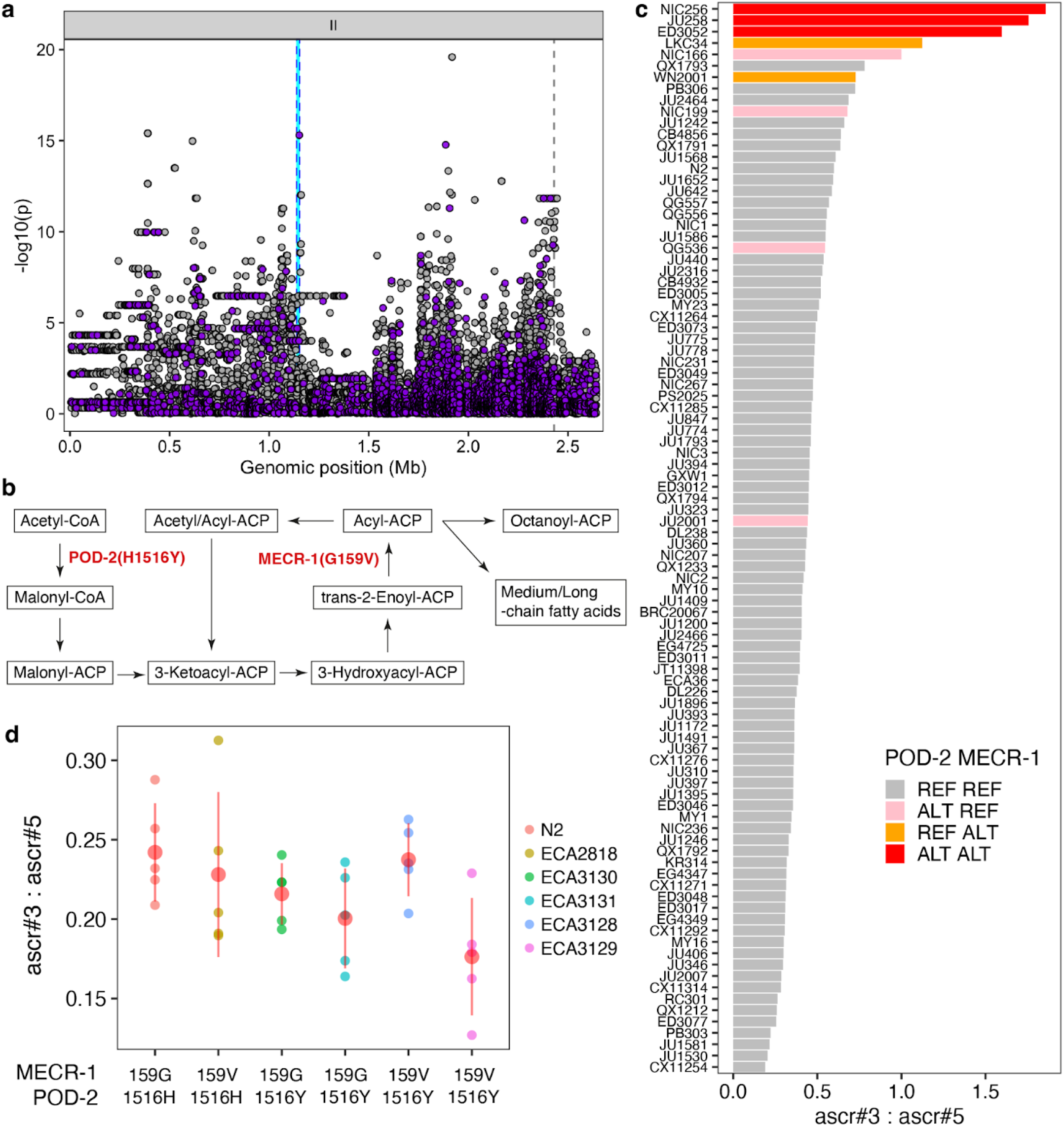
Another mtFAS gene, *pod-2*, is also associated with natural differences in ascaroside production. (a) Fine mapping of the ChrIIL-QTL (II:4,512-2,641,359) for the ascr#3:ascr#5 trait is shown. Each dot represents an SNV that is present in at least 5% of the 94 wild strains. The association between the SNV and ascr#3:ascr#5 trait value is shown on the y-axis and the genomic position of the SNV is shown on the x-axis. SNVs with high or moderate impact inferred from SnpEff are colored purple. (b) Schematic of the mitochondrial FA synthesis (mtFAS) pathway. Two enzymes (POD-2 and MECR-1) that harbor missense variants associated with ascaroside production variation are shown. (c) A bar plot for the ascr#3:ascr#5 trait value of 94 wild *C. elegans* strains. Bars are colored by the genotype of two sites (MECR-1(G159V) and POD-2(H1516Y)). (d) Phenotypes of allele-replaced strains are compared with the N2 reference parental strain (159G, 1516H). On the y-axis, values of ascr#3:ascr#5 ratio traits are shown.

To analyze the effects of the POD-2(H1516Y) variant and its genetic interaction with the MECR-1(G159V) variant, we generated allele substituted strains in both the N2 and the N2 MECR-1(159V) genetic backgrounds. We found that neither the single edit of POD-2(1516H>Y) nor the double edits of both MECR-1(159G>V) and POD-2(1516H>Y) changed the ascr#3:ascr#5 ratio (Fig. 6d). Although these two variants are highly associated with phenotypic variation and components of the same mtFAS pathway, this result shows that neither MECR-1(G159V), POD-2(H1516Y), nor both variants are sufficient to change the ratio of ω-ascarosides to (ω-1)-ascarosides in the N2 background.

## Discussion

We have explored natural variation in ascaroside pheromone production of *C. elegans* and its genetic basis. By profiling excreted metabolites across 95 wild *C. elegans* strains, we found that ascaroside pheromone bouquets differ between strains in several different ways. The most extreme variation was the complete absence of ascaroside pheromones in the JU1400 strain likely caused by a deletion encompassing the peroxisomal β-oxidation gene *daf-22*. Similar to *daf-22(ok693)* laboratory mutants, this wild strain has lost the ability to produce short- and medium-chain ascarosides and instead accumulates long-chain precursors. The natural loss of the *daf-22* gene was surprising because ascaroside pheromones are known to play key roles in the survival and reproduction of the species^4^. Notably, we scanned over 500 wild *C. elegans* genomes and identified this loss only in the genome of the JU1400 strain. This strain was sampled from an urban garden in the city center of Sevilla, Spain, suggesting that this rare *daf-22* deletion has been maintained in a human-associated environment. Similarly, a nonsense mutation in the carboxylesterase *cest-3* in the ECA36 strain was correlated with the lack of indole ascarosides, which regulate dwelling and aggregation behaviors^8^. We also found this *cest-3* variant from the genome of 28 other wild *C. elegans* strains that were sampled across Pacific regions^29^, suggesting that this variant can be maintained in the natural populations.

In addition, our analysis revealed a negative correlation between the relative abundances of ω- and (ω-1)-ascarosides across many different natural strains, highlighted by their most abundant representatives, ascr#5 (ω) and ascr#3 (ω-1). The structural difference between ω- and (ω-1)-ascarosides, which also regulate different phenotypes, likely arises from differences in the metabolism of their long-chain fatty acid precursors, which presumably get hydroxylated in specific positions of the chain followed by attachment of the ascarylose, producing long-chain precursors of either ω- or (ω-1)-ascarosides. Our GWA mapping analysis uncovered hotspot genomic loci that underlie relative abundances of various ascarosides as well as the ascr#3:ascr#5 ratio. Two hotspot QTL on chromosome II were mapped respectively to loci that harbor coding variants in mtFAS pathway genes (*mecr-1* and *pod-2*) that are highly associated with the (ω-1)- to ω-ascaroside ratio. We hypothesize that the mtFAS pathway underlies the balance between the two parallel ascaroside biosynthetic pathways, and genetic variants in the mtFAS pathway contribute to the natural differences in the usage of the two pathways. However, our allele replacement experiments in the N2 strain failed to demonstrate the causal effects of these two variants, which could be interpreted in several ways. First, the mtFAS pathway might not be involved in ascaroside biosynthesis, or at least may not be responsible for the natural variation in (ω-1)- to ω-ascaroside ratio. Second, two variants (MECR-1(G159V), POD-2(H1516Y)) that we edited may be neutral but linked to uncharacterized causal variants in other genes. Finally, complex genetic interactions could mask the effect of these alleles. Recently, incompatible versions of a galactose metabolic pathway were characterized in *Saccharomyces cerevisiae*^*34*^, in which the incompatible combination of alleles of metabolic genes is not found in nature. We failed to introduce MECR-1(159G>V) edit in two wild strains while successfully introducing MECR-1(159V>G) edit in the N2 background, implying that incompatible alleles of metabolic genes might be present across wild genomes of *C. elegans*. Notably, we found strong LD among three ascr#3:ascr#5 QTL (the *mecr-1* locus, the *pod-2* locus, and the ChrIV-QTL). Therefore, although even the double edits (MECR-1(159G>V) and POD-2(1516H>Y)) did not display effects on the mapped trait (ascr#3:ascr#5), this result could be explained by allele(s) in unidentified genes that segregate together and exert non-additive phenotypic effects.

Although we focused on the (ω-1)-ascaroside to ω-ascaroside ratio among many observed traits in this study, we also discovered natural variation in the production of individual ascaroside pheromones and their unique QTL. Our dataset will provide a valuable resource for future studies to characterize novel genes involved in pheromone production and to explore the molecular mechanisms of how genetic changes lead to the evolution of a chemical language.

## Methods

### *C. elegans* strains and growth

N2 (Bristol) and wild nematode strains were maintained at 20 °C reared on *E. coli* OP50 and grown on modified nematode growth medium containing 1% agar and 0.7% agarose (NGMA) to prevent animals from burrowing^35^. Wild strains were obtained from the CeNDR^29^ and are available upon request. Strain information can be found in the CeNDR. For the analysis of staged adults, approximately 35,000 synchronized L1 larvae obtained from alkaline bleach treatment were added to 125 mL Erlenmeyer flasks containing S-Complete medium at a density of ∼3,000 worms / mL. Nematodes were fed with concentrated *E. coli* OP50 and incubated at 20 °C with shaking at 180 RPM for approximately 64 – 70 h, at which time the population was predominantly gravid adults as determined by microscopic inspection. Liquid cultures were transferred to 15 mL conical tubes and centrifuged (500 x g, 22 °C, 1 min), and the supernatant (conditioned media, *exo*-metabolome) was transferred to a fresh conical tube and snap frozen.

### Sample preparation

*Exo*-metabolome (conditioned media) samples were lyophilized for 24 hrs using a VirTis BenchTop 4 K Freeze Dryer. Dried material was directly extracted in 3 mL of methanol with gentle rocking at room temperature. Following overnight extraction, samples were centrifuged (2,750 x g, 22 °C, 5 min) in an Eppendorf 5702 Centrifuge. The supernatant was transferred to a clean 8 mL glass vial and concentrated to dryness in an SC250EXP Speedvac Concentrator coupled to an RVT5105 Refrigerated Vapor Trap (Thermo Scientific). The powder was suspended in 150 μL of methanol, vortexed vigorously for 30 seconds and sonicated for five minutes. The suspension was transferred to a 1.7 mL Eppendorf tube and centrifuged (18,000 x g, 22 °C, 5 min), and the clarified supernatant was transferred to HPLC vials and analyzed directly by HPLC-HRMS, see below.

### High-performance liquid chromatography coupled to high-resolution mass spectrometry

Liquid chromatography was performed on a Vanquish HPLC system controlled by Chromeleon Software (ThermoFisher Scientific) and coupled to an Orbitrap Q-Exactive High Field mass spectrometer controlled by Xcalibur software (ThermoFisher Scientific). Methanolic extracts prepared as described above were separated on a Thermo Hypersil Gold C18 column (150 mm × 2.1 mm, particle size 1.9 μM, part no. 25002-152130) maintained at 40 °C with a flow rate of 0.5 mL/min. Solvent A: 0.1% formic acid (Fisher Chemical Optima LC/MS grade; A11750) in water (Fisher Chemical Optima LC/MS grade; W6-4); solvent B: 0.1% formic acid in acetonitrile (Fisher Chemical Optima LC/MS grade; A955-4). A/B gradient started at 1% B for 3 min after injection and increased linearly to 98% B at 20 min, followed by 5 min at 98% B, then back to 1% B over 0.1 min and finally held at 1% B for the remaining 2.9 min to re-equilibrate the column (28 min total method time). Mass spectrometer parameters: spray voltage, −3.0 kV/+3.5 kV; capillary temperature 380 °C; probe heater temperature 400 °C; sheath, auxiliary, and sweep gas, 60, 20, and 2 AU, respectively; S-Lens RF level, 50; resolution, 120,000 at *m/z* 200; AGC target, 3E6. Each sample was analyzed in negative (ESI−) and positive (ESI+) electrospray ionization modes with *m/z* range 100–1000.

### Identification of a large deletion at *daf-22* locus in JU1400

To identify structural variants in JU1400, we downloaded a genome assembly for JU1400 assembled with PacBio long reads^36^ from NCBI (GCA_016989365.1) and called structural variants using MUM&Co^37^ (version 3.8; default parameters) with the N2 genome (WS285) as a reference. MUM&Co identified a 29,011 bp deletion (II:12411041-12440052) in JU1400 relative to N2 that overlaps with *daf-22* and seven other protein coding genes (*Y57A10C*.*1, Y57A10C*.*11, sre-27, sre-26, Y57A10C*.*8, dct-12, Y57A10C*.*9*). We confirmed this deletion by aligning unassembled PacBio long reads for JU1400 (PRJNA692613) to the N2 reference genome using minimap2^38^ (version 2.17; using the parameters *-a -x map-pb*) and inspecting read coverage using IGV^39^ (version 2.8.13).

### Genetic relatedness

A VCF file containing 963,027 biallelic SNVs from a previous study^36^ was filtered for 94 wild *C. elegans* strains and converted to the PHYLIP format. The distance matrix and pseudo-rooted (ECA36) neighbor-joining tree were made from this PHYLIP file using dist.ml and the NJ function using the phangorn (version 2.5.5) R package. The tree was visualized using the ggtree (version 1.16.6) R package.

### Heritability calculations

Narrow-sense heritability (*h*^2^) estimates were calculated using the phenotype data of 94 wild strains. The *A*.*mat* functions in the sommer R package^40^ were used to generate an additive genotype matrix, from the genotype matrix used for the GWA mapping. This matrix was used to calculate the additive variance components using the sommer *mmer* function. Variance components were used to estimate heritability and standard error through the *pin* function (*h*^*2*^ ∼ V1 / V1 + V2) in the sommer package.

### Principal component analysis

Phenotypic values for heritable (>10% narrow-sense heritability) relative abundance traits of 23 ascarosides were used as inputs to principal component analysis (PCA). PCA was performed using the prcomp function in R. Eigenvectors and loadings were subsequently extracted from the object returned by the prcomp function.

### GWA mapping

A genome-wide association mapping was performed for heritable (>10%) relative abundance traits of 23 ascarosides as well as the ascr#3 to ascr#5 (ascr#3:ascr#5) ratio trait using the NemaScan pipeline^41^ available at https://github.com/AndersenLab/NemaScan. Genotype data were acquired from the latest VCF release (release 20210121) from the CeNDR. We used BCFtools^42^ to filter variants below a 5% minor allele frequency and variants with missing genotypes and used PLINK v1.9^43,44^ to prune genotypes using linkage disequilibrium. The additive kinship matrix was generated from the 30,065 markers using the *make-grm* and *make-grm-inbred* functions from GCTA^45^. Because these markers have high LD, we performed eigen decomposition of the correlation matrix of the genotype matrix to identify 499 independent tests^46^.We performed genome-wide association mapping using the *mlma-loco* and *fastGWA-lmm-exact* functions from GCTA. Significance was determined by an eigenvalue threshold set by the number of independent tests in the genotype matrix^46^. Confidence intervals were defined as +/- 150 SNVs from the rightmost and leftmost markers that passed the significance threshold.

### CRISPR-Cas9 allele replacement

Genome editing to make alleles *pod-2(ean229), pod-2(ean238), pod-2(ean239), pod-2(ean240), mecr-1(ean216), mecr-1(ean220)* was done using CRISPR-Cas9 and the co-conversion marker *dpy-10* as previously described^47^. Single-strand guide RNAs (sgRNAs) for *mecr-1* and *pod-2* were designed using the online analysis platform Benchling (benchling.com). All guides were ordered from Synthego (Redwood City, CA) and injected at 1 μM for the *dpy-10* guide and 6 μM for all others. Single-stranded oligodeoxynucleotides (ssODN) templates used for homology-directed repair (HDR) were ordered from Integrated DNA and injected at 0.5 μM for the *dpy-10* ssODN and 5 μM for all others, and 5 μM of purified Cas9 protein (QB3 Macrolab, Berkeley, CA) was used. Hermaphrodites were staged at L4 larval stage the day before injection, and the reagents were mixed and incubated for one hour at room temperature prior to injection into each gonad. Injected animals were singled onto NGMA plates and allowed to lay until the next generation matured to the L4/young adult stage. Plates were screened for the *dpy-10* phenotypes of Dumpy and Roller and F1s were singled from plates with a high percentage of affected worms. F1s were allowed to lay eggs before single animal lysis and PCR, and the products were sequenced using Sanger sequencing by MCLab Molecular Cloning Laboratories (South San Francisco, CA) with no more than one edited strain per independently injected progenitor retained for study. All alleles were confirmed by sequencing singled offspring for at least two additional generations to confirm the accuracy of the edit sequence and the homozygosity of the line.

All genome edited strains through CRISPR are listed below:

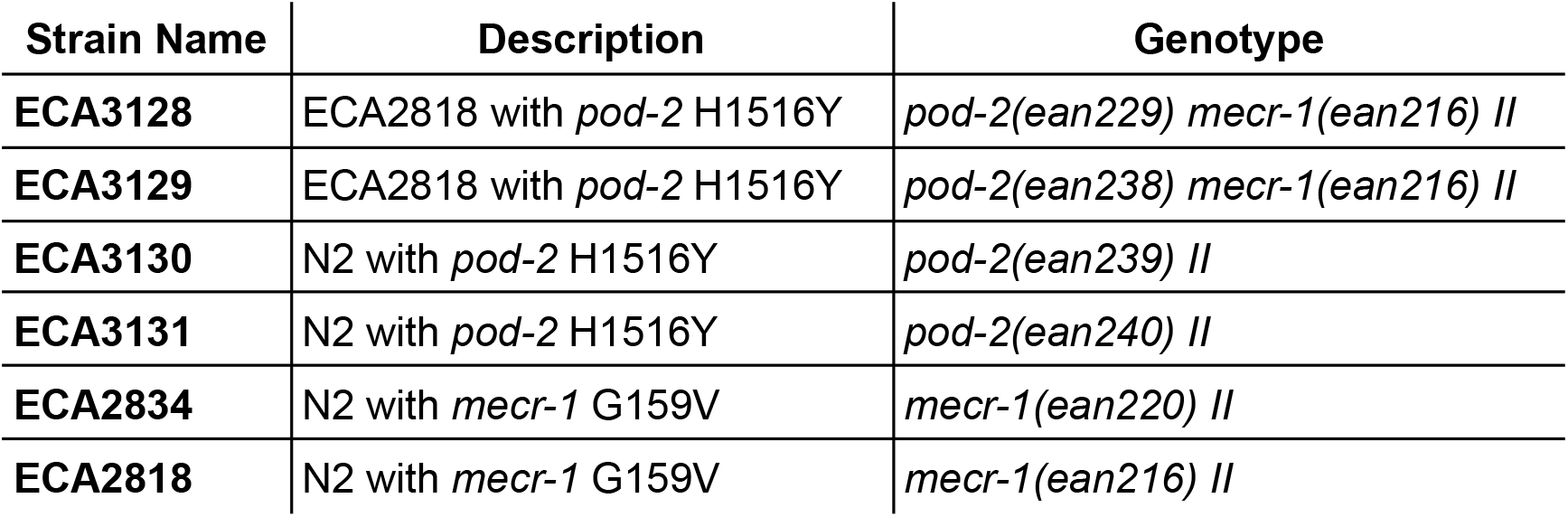

Oligonucleotides used for the generation of genome-edited strains are listed below:

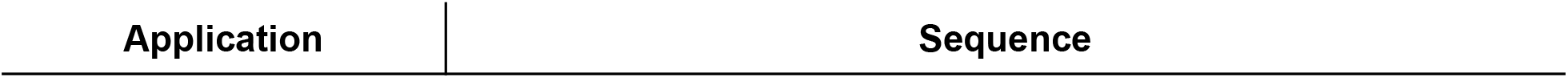
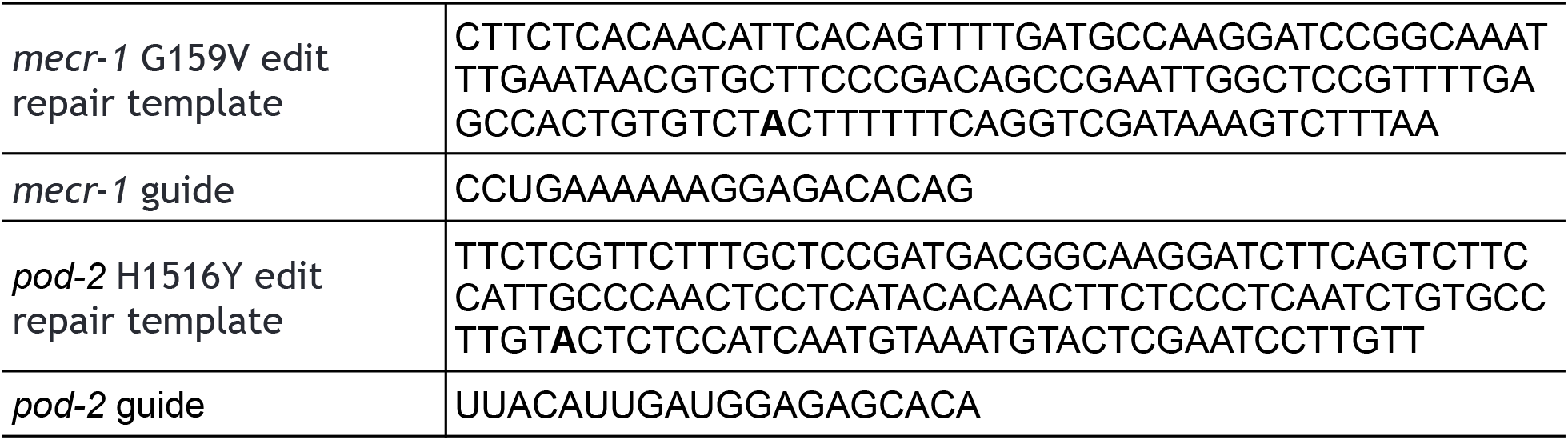

## Data and code availability

All data sets and code for generating figures and tables are available on GitHub (https://github.com/AndersenLab/Ce-ascr).

## Acknowledgments

This work was funded by an NSF CAREER award (1751035) to E.C.A. and grants from the National Institutes of Health (R35 GM131877-05 to F.C.S. and DK115690 to E.C.A. and F.C.S.). The *Caenorhabditis* Natural Diversity Resource is supported by a National Science Foundation Living Collections Award to E.C.A. (NSF Capacity grant 2224885). Some strains were provided by the CGC, which is funded by NIH Office of Research Infrastructure Programs (P40 OD010440).

## Author contributions

D.L., F.C.S., and E.C.A. conceived and designed the study. D.L., B.W.F., F.C.S., and E.C.A. analyzed the data and wrote the manuscript. O.P., D.C.F.P., F.J.T., B.W.F., P.R.R., and A.R.K. prepared cultures and/or performed HPLC-HRMS analyses. E.J.K. performed CRISPR-Cas9 allele replacement experiments. D.L. and K.S.E. performed genome-wide association mapping analysis. L.S. identified deletion in *daf-22* locus of JU1400.

## Additional information

**Supplementary information** is available for this paper.

**Correspondence and requests for materials** should be addressed to E.C.A. and F.C.S.

