## Supplementary Information for "Natural genetic variation in the pheromone production of *C. elegans*"

Daehan Lee, Bennett W. Fox, Diana C. F. Palomino, Oishika Panda, Francisco J. Tenjo, Emily J. Koury, Kathryn S. Evans, Lewis Steven, Pedro R. Rodrigues, Aiden R. Kolodziej, Frank C. Schroeder*, Erik C. Andersen*

**Supplementary Fig. 1 | Structures of 44 ascarosides**

Chemical structures of the 44 ascarosides included in the analysis.

### “Simple” ascarosides

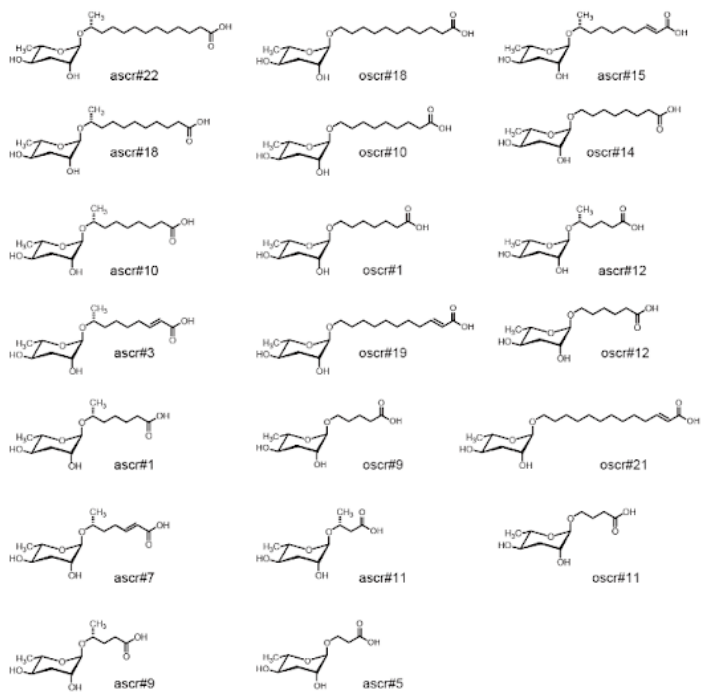


### b-hydroxy ascarosides

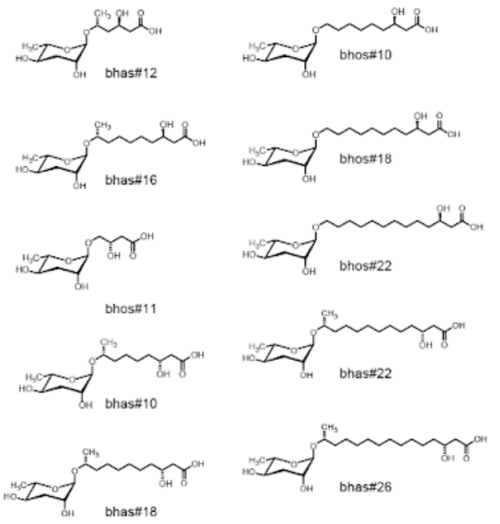


### *C*-term and 4’ modified ascarosides

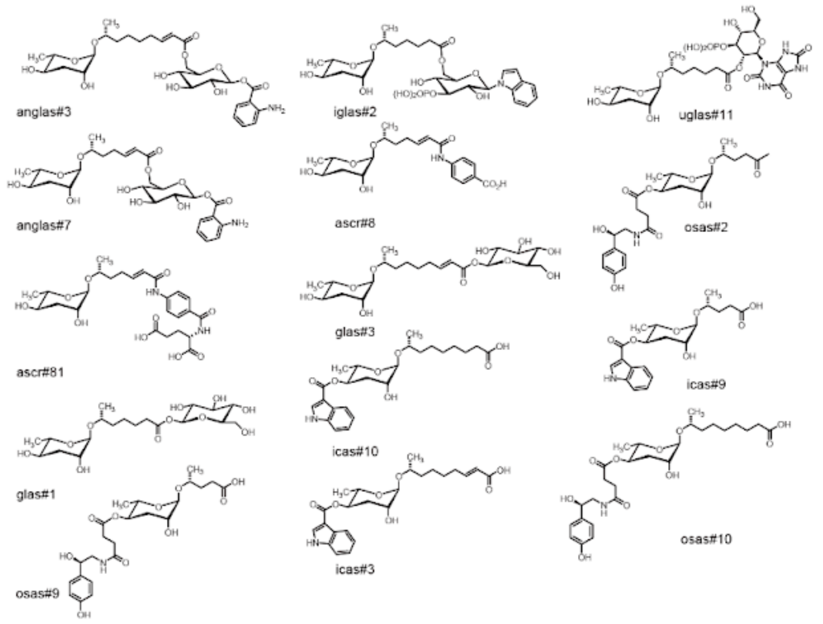

**Supplementary Fig. 2 | Natural variation in the abundances of 42 ascaroside compounds**
Bar plots showing relative abundances of 42 ascaroside compounds across 94 wild *C. elegans* strains, ordered by the relative abundance of each trait.

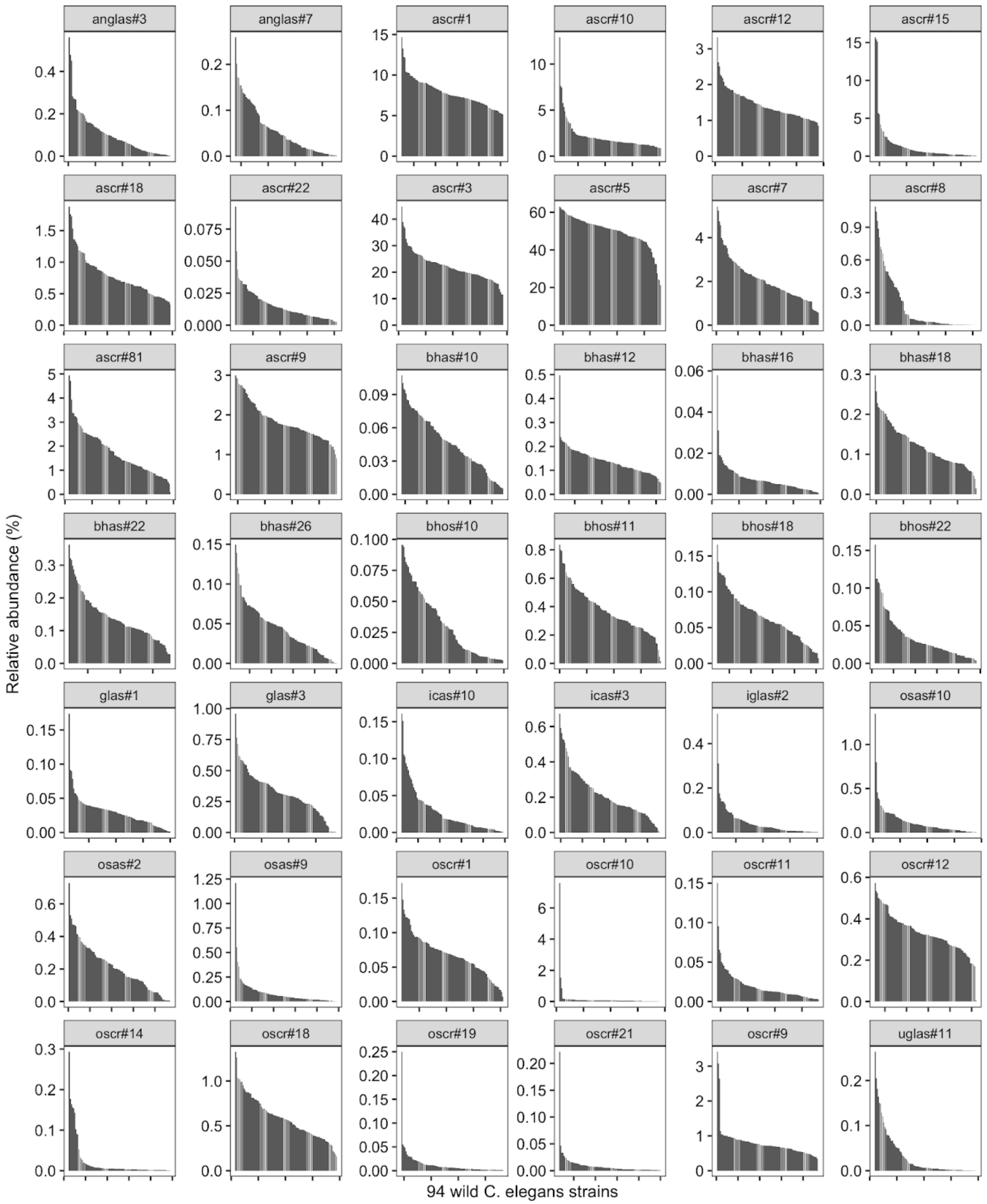

**Supplementary Fig. 3 | Phenotypes of MECR-1 G159V allele-replacement strains**

Phenotypes of MECR-1(G159V) allele-replaced strains are compared with the N2 reference parental strain (159G) and two wild strains with MECR-1(159V). On the y-axis, the relative ratios between ascr#3 and ascr#5 are shown.

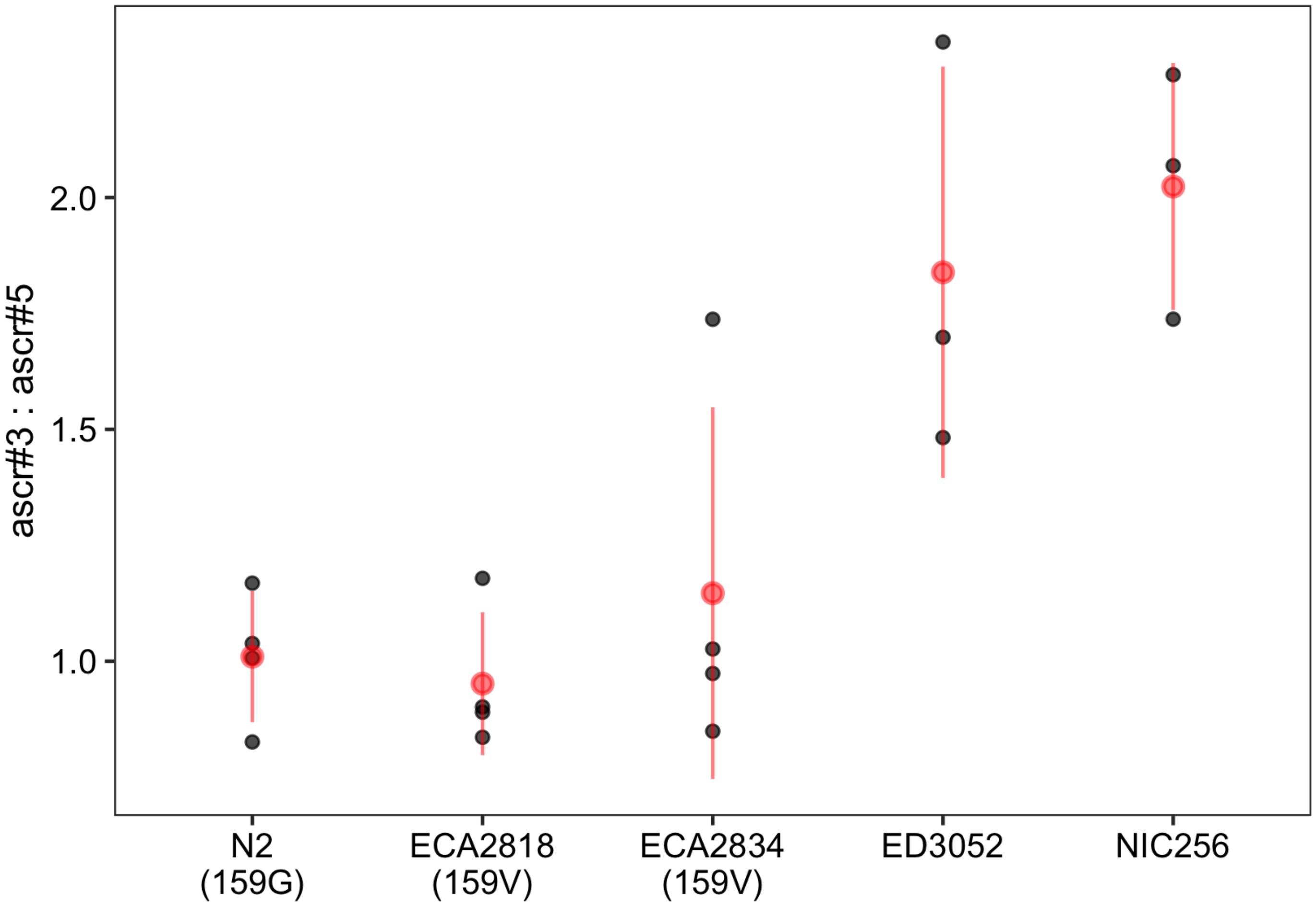

**Supplementary Fig. 4 | Linkage disequilibrium among ascr#3:ascr#5 QTL**
Linkage disequilibrium ($r^2$) values of four peak QTL markers for ascr#3:ascr#5 trait are shown.

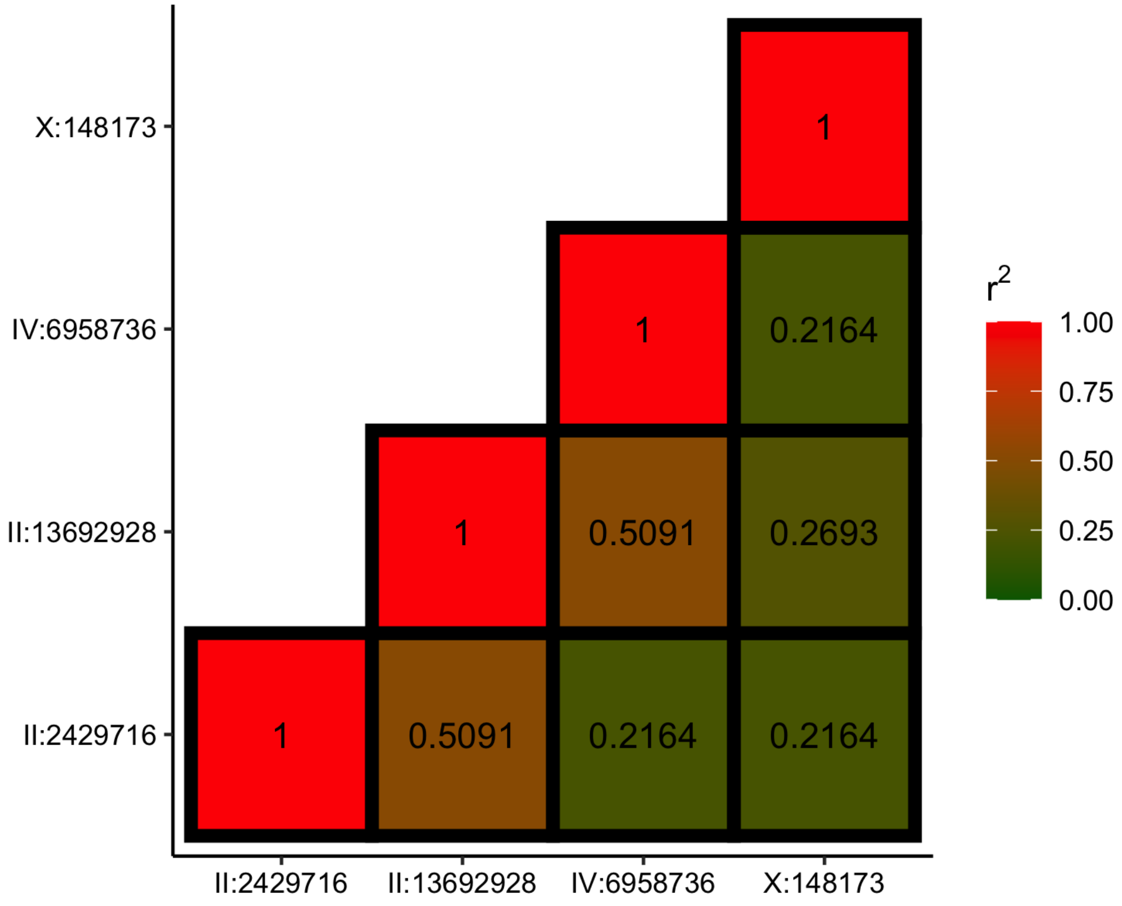

**Supplementary Fig. 5 | Phenotypic variance explained by the POD-2(H1516Y) variant**
A heatmap showing amounts of variance explained by the POD-2(H1516Y) variant for pairwise ratio traits of 23 ascarosides with high heritability.

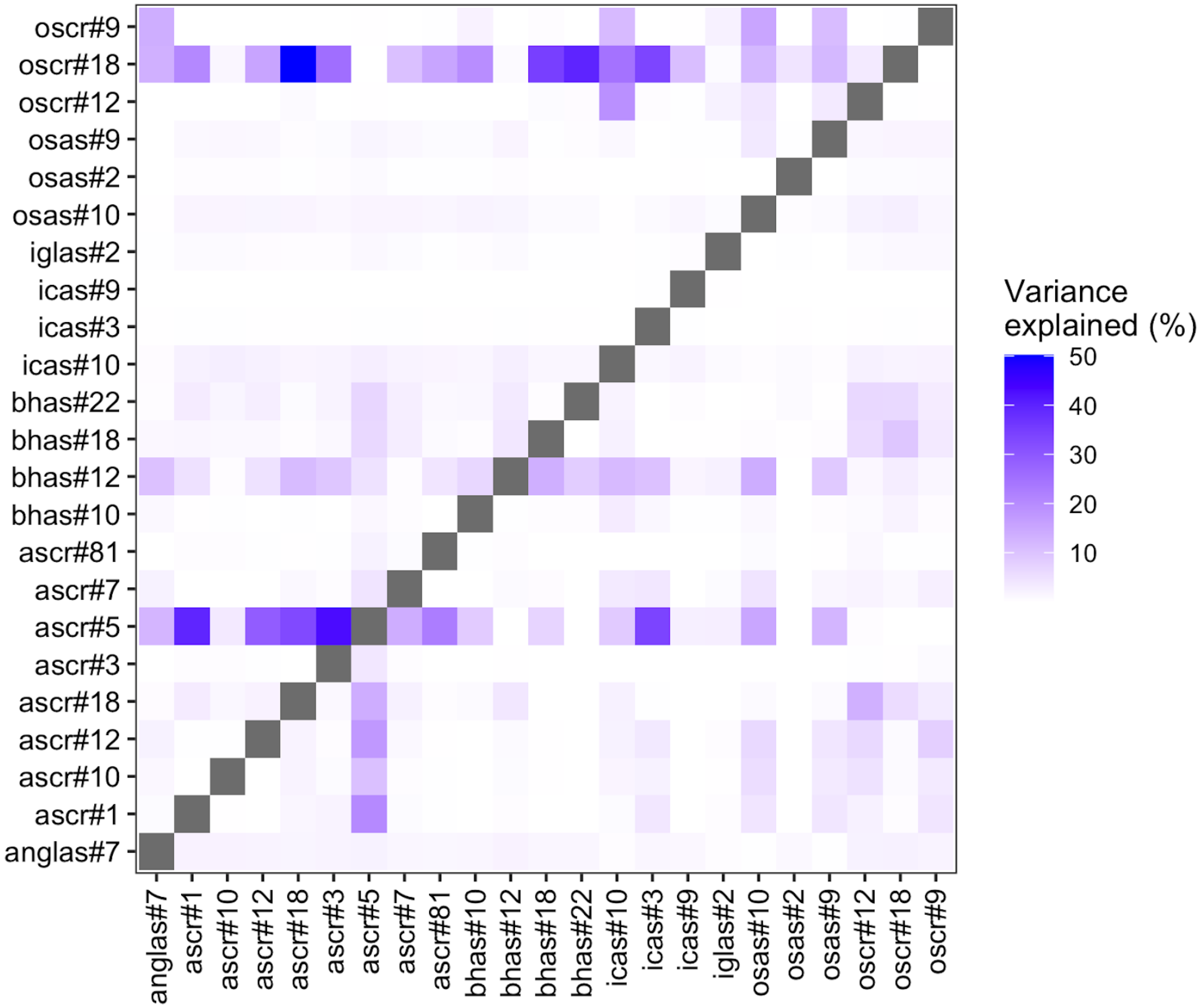

**Supplementary Table 1 | Summary of GWA mapping**

| Trait | Chr | Start | Peak | End | Interval size (bp) | Variance explained | Log10p |
|---|---|---|---|---|---|---|---|
| anglas#7 | I | 3736 | 2733130 | 14747808 | 14744072 | 0.356921053 | 12.04815028 |
| anglas#7 | II | 4512 | 1093248 | 6846802 | 6842290 | 0.228572941 | 6.749824052 |
| anglas#7 | II | 13399659 | 14075021 | 14893958 | 1494299 | 0.19468924 | 5.619052891 |
| anglas#7 | III | 373310 | 2125160 | 3787190 | 3413880 | 0.225667053 | 6.649273358 |
| anglas#7 | IV | 4272318 | 13598850 | 17489019 | 13216701 | 0.268041954 | 8.189610868 |
| anglas#7 | V | 593381 | 13756973 | 20654084 | 20060703 | 0.291770096 | 9.128340681 |
| anglas#7 | X | 124570 | 2287865 | 8740424 | 8615854 | 0.336113477 | 11.05528198 |
| anglas#7 | X | 10742782 | 15832204 | 16115328 | 5372546 | 0.212987172 | 6.21857424 |
| ascr#1 | II | 13198273 | 13390616 | 13852723 | 654450 | 0.199305394 | 5.767942064 |
| ascr#1 | III | 11741084 | 11994388 | 12747249 | 1006165 | 0.218405445 | 6.401035591 |
| ascr#1 | V | 17709799 | 17979776 | 18414183 | 704384 | 0.222195319 | 6.53005581 |
| ascr#1 | X | 1268287 | 2334653 | 7798884 | 6530597 | 0.223316347 | 6.568443259 |
| ascr#10 | I | 92616 | 11607099 | 12296778 | 12204162 | 0.262834831 | 7.752245163 |
| ascr#10 | III | 176496 | 610190 | 856556 | 680060 | 0.195658939 | 5.48507788 |
| ascr#10 | X | 124570 | 1796524 | 8206669 | 8082099 | 0.247626226 | 7.205905297 |
| ascr#12 | II | 12829939 | 13412550 | 13830067 | 1000128 | 0.187873388 | 5.402014285 |
| ascr#12 | X | 3675721 | 5654087 | 6514350 | 2838629 | 0.187851903 | 5.401335315 |
| ascr#15 | X | 13834552 | 14603523 | 16845496 | 3010944 | 0.210973885 | 5.970478244 |
| ascr#22 | IV | 43186 | 773445 | 1056922 | 1013736 | 0.225775256 | 6.587458817 |
| ascr#3 | II | 303114 | 2429716 | 2641359 | 2338245 | 0.310848244 | 6.01691297 |
| ascr#3 | II | 12476063 | 13203646 | 13886985 | 1410922 | 0.425392871 | 8.688487716 |
| ascr#5 | I | 695386 | 1233659 | 2260201 | 1564815 | 0.212354447 | 6.197417817 |
| ascr#5 | II | 4512 | 786255 | 2641359 | 2636847 | 0.235902331 | 7.006610949 |
| ascr#5 | II | 12368778 | 13692928 | 15278446 | 2909668 | 0.433081098 | 16.28871339 |
| ascr#5 | III | 11479501 | 11775202 | 12781165 | 1301664 | 0.253949117 | 7.659026725 |
| ascr#5 | IV | 751766 | 13489747 | 15131605 | 14379839 | 0.232898852 | 6.900830594 |
| ascr#5 | X | 124570 | 2986710 | 6700584 | 6576014 | 0.243249564 | 7.268690917 |
| ascr#5 | X | 10603327 | 12636855 | 13504734 | 2901407 | 0.195723586 | 5.652279783 |
| ascr#7 | II | 2739775 | 2867991 | 3096129 | 356354 | 0.212968229 | 6.217940367 |
| ascr#7 | II | 12368778 | 12716976 | 13298659 | 929881 | 0.203992488 | 5.920724667 |
| ascr#81 | II | 4013239 | 4492510 | 6846802 | 2833563 | 0.188183858 | 5.411829173 |

| | | | | | | | |
|---|---|---|---|---|---|---|---|
| ascr#81 | II | 13298548 | 15081782 | 15278446 | 1979898 | 0.203178 | 5.894057439 |
| ascr#81 | III | 11727315 | 12525622 | 13323719 | 1596404 | 0.240632753 | 7.17480245 |
| ascr#81 | IV | 3981501 | 13598850 | 15830951 | 11849450 | 0.376011609 | 13.01582251 |
| ascr#81 | V | 3852565 | 12897665 | 20909920 | 17057355 | 0.230052293 | 6.801287943 |
| ascr#81 | X | 124570 | 2287865 | 8468772 | 8344202 | 0.278411575 | 8.592593798 |
| ascr#81 | X | 10402729 | 12898474 | 14134472 | 3731743 | 0.205360013 | 5.965612216 |
| ascr#9 | X | 124570 | 867999 | 8196736 | 8072166 | 0.201037248 | 5.824204162 |
| bhas#12 | III | 11479501 | 11801670 | 13775378 | 2295877 | 0.197417124 | 5.706851883 |
| bhas#12 | V | 639357 | 1648854 | 2121953 | 1482596 | 0.193271526 | 5.573636782 |
| bhas#18 | II | 4512 | 1304965 | 2641359 | 2636847 | 0.20995443 | 6.117453264 |
| bhas#18 | II | 12542769 | 13692928 | 13886985 | 1344216 | 0.268616618 | 8.21165837 |
| bhas#18 | IV | 1784821 | 3579831 | 13994964 | 12210143 | 0.197750073 | 5.717604495 |
| bhas#18 | X | 10644948 | 11384708 | 13555358 | 2910410 | 0.208321824 | 6.063311369 |
| bhas#22 | II | 4512 | 1304965 | 1819726 | 1815214 | 0.196040546 | 5.662478844 |
| bhas#22 | II | 12542769 | 13692928 | 13852723 | 1309954 | 0.219786155 | 6.447905139 |
| bhas#22 | IV | 1670573 | 2053138 | 12976479 | 11305906 | 0.244592202 | 7.317100755 |
| bhos#22 | IV | 1489441 | 1658768 | 4903773 | 3414332 | 0.206928314 | 6.017260795 |
| bhos#22 | V | 16774869 | 16912293 | 17723744 | 948875 | 0.190747949 | 5.493148752 |
| icas#10 | X | 12993889 | 13280531 | 14746716 | 1752827 | 0.324310354 | 7.034667983 |
| icas#3 | IV | 1458400 | 2153686 | 17187855 | 15729455 | 0.259176934 | 7.853617666 |
| icas#3 | X | 4114709 | 5020050 | 5834064 | 1719355 | 0.191589274 | 5.519931225 |
| icas#3 | X | 9782566 | 11067399 | 12958823 | 3176257 | 0.243679952 | 7.284191297 |
| icas#9 | II | 4512 | 287101 | 930896 | 926384 | 0.23459298 | 6.822812808 |
| icas#9 | II | 3376344 | 3805135 | 5699108 | 2322764 | 0.205637608 | 5.857879705 |
| icas#9 | IV | 1020003 | 1196322 | 3699373 | 2679370 | 0.265917324 | 7.946541235 |
| icas#9 | X | 4099789 | 11067399 | 12388111 | 8288322 | 0.278051744 | 8.406627526 |
| osas#10 | IV | 1489441 | 15194433 | 16785392 | 15295951 | 0.220790754 | 6.354586574 |
| osas#10 | X | 1499830 | 8910902 | 15347272 | 13847442 | 0.256054219 | 7.583086688 |
| osas#2 | I | 94720 | 3951912 | 13659314 | 13564594 | 0.259253902 | 7.856501184 |
| osas#2 | II | 11998593 | 13709164 | 14893958 | 2895365 | 0.227210457 | 6.702591881 |
| osas#2 | III | 10477290 | 11650544 | 13464737 | 2987447 | 0.240371153 | 7.1654496 |
| osas#2 | IV | 13180663 | 15550884 | 16120945 | 2940282 | 0.217812207 | 6.380944254 |
| osas#2 | V | 1583467 | 13965981 | 19804412 | 18220945 | 0.213860232 | 6.247817825 |
| osas#2 | X | 388657 | 2362150 | 8740424 | 8351767 | 0.244905162 | 7.328407881 |
| osas#2 | X | 12921258 | 14341555 | 16241564 | 3320306 | 0.195231967 | 5.636478272 |

| | | | | | | | |
|---|---|---|---|---|---|---|---|
| osas#9 | IV | 9103306 | 15559488 | 16820852 | 7717546 | 0.214502498 | 6.146323686 |
| osas#9 | X | 124570 | 4960069 | 17379408 | 17254838 | 0.227130596 | 6.567728384 |
| oscr#12 | II | 12985307 | 13204800 | 13597625 | 612318 | 0.229014473 | 6.765166475 |
| oscr#12 | III | 11507368 | 11801670 | 12028729 | 521361 | 0.193085092 | 5.567675418 |
| oscr#14 | X | 13179717 | 13882309 | 14622118 | 1442401 | 0.228300473 | 6.074616735 |
| oscr#18 | I | 1738955 | 12959042 | 15064788 | 13325833 | 0.261215584 | 7.930206146 |
| oscr#19 | X | 16309679 | 16879447 | 17536376 | 1226697 | 0.191635451 | 5.467732405 |
| oscr#21 | II | 12910581 | 13692928 | 13852955 | 942374 | 0.199129391 | 5.706045529 |
| oscr#9 | I | 7830004 | 12302651 | 13784685 | 5954681 | 0.229639502 | 6.585974782 |
| oscr#9 | III | 11816876 | 12374645 | 13775378 | 1958502 | 0.20371408 | 5.73826383 |
| uglas#11 | II | 13175817 | 13358149 | 13705977 | 530160 | 0.200120674 | 5.737871785 |
| uglas#11 | X | 14477147 | 15104579 | 15590617 | 1113470 | 0.200843107 | 5.76111123 |
| ascr#3: ascr#5 | II | 4512 | 2429716 | 2641359 | 2636847 | 0.360079666 | 6.831116714 |
| ascr#3: ascr#5 | II | 12422412 | 13692928 | 13915254 | 1492842 | 0.717846993 | 14.2504631 |
| ascr#3: ascr#5 | IV | 4212020 | 6958736 | 10752480 | 6540460 | 0.450031818 | 6.655576285 |
| ascr#3: ascr#5 | X | 124570 | 148173 | 4987366 | 4862796 | 0.356242553 | 6.169168913 |
